# Natural variation in codon bias and mRNA folding strength interact synergistically to modify protein expression in *Saccharomyces cerevisiae*

**DOI:** 10.1101/2022.09.07.507001

**Authors:** Anastacia N. Wienecke, Margaret L. Barry, Daniel A. Pollard

## Abstract

Codon bias and mRNA folding strength (mF) are hypothesized molecular mechanisms by which polymorphisms in genes modify protein expression. Natural patterns of codon bias and mF across genes as well as effects of altering codon bias and mF suggest the influence of these two mechanisms may vary depending on the specific location of polymorphisms within a transcript. Despite the central role codon bias and mF may play in natural trait variation within populations, systematic studies of how polymorphic codon bias and mF relate to protein expression variation are lacking. To address this need, we analyzed genomic, transcriptomic, and proteomic data for 22 *Saccharomyces cerevisiae* isolates, estimated protein accumulation for each allele of 1620 genes as the log of protein molecules per RNA molecule (logPPR), and built linear mixed effects models associating allelic variation in codon bias and mF with allelic variation in logPPR. We found codon bias and mF interact synergistically in a positive association with logPPR and this interaction explains almost all the effect of codon bias and mF. We examined how the locations of polymorphisms within transcripts influence their effects and found that codon bias primarily acts through polymorphisms in domain encoding and 3’ coding sequences while mF acts most significantly through coding sequences with weaker effects from UTRs. Our results present the most comprehensive characterization to date of how polymorphisms in transcripts influence protein expression.

## Introduction

Decades of research efforts have established that heritable variation in protein expression is a major driver of higher-order trait variation (Chan et al., 2010; Skelly et al., 2009; Stern and Orgogozo, 2008). Advances in nucleic acid quantification technologies have facilitated numerous studies probing the effects of molecular polymorphisms on mRNA abundance variation (Brem et al., 2002; Pai et al., 2012; Rockman and Kruglyak, 2006). This work established that the genetic architecture of gene expression is divided into two parts: a modest number of polymorphisms act in *trans* on the expression of many genes, and a large number act allele-specifically in *cis*. More recent studies have focused on protein abundances and found that genetic variation commonly acts specifically at the protein level, modifying either protein synthesis or decay rates (Albert et al., 2014; Foss et al., 2011; Gygi et al., 1999; Parts et al., 2014; Pollard et al., 2016; Straub, 2011; Torabi and Kruglyak, 2011). Despite enormous progress establishing that polymorphisms act in both *cis* and *trans* as well as at the mRNA-level and protein-level, the diversity of molecular mechanisms by which polymorphisms act on protein expression abundances remains poorly resolved (Courtier-Orgogozo et al., 2020; Nieuwkoop et al., 2020).

Codon bias and mRNA folding stability (mF) are two hypothesized mechanisms by which polymorphisms act in *cis* on protein expression (Hanson and Coller, 2018; Tuller et al., 2011). Both mechanisms have been studied using various approaches. This includes comparing the codon bias and mF of different genes within a genome (Dana and Tuller, 2014; Zur and Tuller, 2012), comparing them between species (LaBella et al., 2019; Park et al., 2013), engineering alleles with artificially modified codon bias and mF (Babendure et al., 2006; Gooch et al., 2008), and computationally modeling their impact on protein expression (Mao et al., 2014; Tuller et al., 2011).

To our knowledge, no study has systematically investigated how natural polymorphic variation in codon bias and mF relate to variation in protein expression. Investigating these factors in a population context is important for several reasons. Comparisons amongst alleles of the same gene, instead of comparisons across genes within the genome of an individual, minimizes potential confounding effects. Because standing allelic variation is typically comprised of complex combinations of genetic differences, population studies can reveal effects that are distinct from those seen from traditional single perturbation mutagenesis experiments (Greenspan, 2004). Furthermore, population studies have direct relevance for understanding human population variation and for the broader goal of characterizing molecular evolutionary mechanisms.

Uneven synonymous codon usage is referred to as codon bias and the overall pattern of codon bias in a species’ genome is understood to be the result of two factors (Hershberg and Petrov, 2008; LaBella et al., 2019; Plotkin and Kudla, 2011; Trotta, 2013; Wallace et al., 2013). First, species-specific mutational biases produce codons at different rates; yeast DNA, for example, mutates to AT nucleotides at approximately twice the rate as GC nucleotides (Lynch et al., 2008). Second, natural selection appears to favor specific codons over others (LaBella et al., 2019). For instance, codon usage in highly expressed genes is unique relative to that of the whole genome. Their most frequently used codons tend to be those with high abundances of cognate tRNAs, presumably because the ribosome translates these codons fastest and with the most precision (Dana and Tuller, 2014; Ikemura, 1982; LaBella et al., 2019; Sharp et al., 1986); we refer to these codons as translationally optimal codons. This mechanistic model is supported by the observed upregulation of genes with codons well matched to a characteristic fluctuation in tRNA supply (Quax et al., 2015) (e.g. as occurs during the cell-cycle (Frenkel-Morgenstern et al., 2012), circadian rhythms (Xu et al., 2013), cell proliferation/differentiation (Gingold et al., 2014), and stress (Torrent et al., 2018)).

Protein abundances can be altered by engineering genes with either favored or unfavored codons (Burgess-Brown et al., 2008; Gooch et al., 2008), however, the impact of polymorphisms that alter codon bias in natural populations remains unexplored. We expect that polymorphisms that increase codon bias would, on average, result in alleles that are more quickly synthesized into protein (see Table 1).

**Table 1.** Predicted association between codon bias and protein expression across alleles.

|  |  |
| --- | --- |
| Region | Predicted association |
| Whole CDS | Positive |
| 5' Region of CDS | None |
| Domain Regions | Positive |
| Inter-Domain Linker Regions | None |
| 3' Region of CDS | Positive |

Codons with lowly abundant cognate tRNAs, which we refer to as translationally suboptimal codons, are most commonly found in the 5’ coding region and in the regions encoding inter-domain linkers (Tuller et al., 2010, 2011; Weinberg et al., 2016). The first 30-50 codons of mRNA, the 5’ coding region, harbor a high density of ribosomes - nearly three times that of any other mRNA region (Ingolia et al., 2009). This pattern is attributed to selection that either slows translation initiation or spreads-out ribosomes such that sufficient spacing between ribosomes avoids downstream collisions and traffic jams that can result in premature translation termination and lower protein synthesis rates (Chu et al., 2014; Doma and Parker, 2006; Tuller et al., 2010). The inter-domain linkers lie in-between protein domains and are some of the most mildly structured protein regions. The slow translation of these areas could facilitate the proper co-translational folding of preceding protein domains, maintaining high levels of stable protein (Makhoul and Trifonov, 2002; Pechmann and Frydman, 2013; Thanaraj and Argos, 1996). We hypothesize that the maintenance of these patterns would constrain selection for high codon bias in these regions. Thus, amongst the alleles of a gene in a natural population, we expect to see a weak association between translation rates and codon bias in these regions (see Table 1).

Translationally optimal codons are typically found in regions encoding protein domains and in the 3’ coding region. This 3’coding region is bordered by the 3’-most domain-encoding region and the translation stop codon. It has the highest proportion of optimal codons of all regions of the CDS (Tuller et al., 2010). Such levels of bias are thought to protect against ribosome collisions and the ensuing interruptions in protein synthesis, such as premature translation termination. It is especially costly in terms of expended energy and resources if a ribosome terminates prematurely this far past the start codon (Plotkin and Kudla, 2011; Tuller et al., 2010). Domain-encoding regions show this pattern presumably because selection for high codon bias is unconstrained and perhaps because selection to maintain domain function additionally selects for the high bias codons that tend to be translated more accurately (Drummond and Wilke, 2009; Geiler-Samerotte et al., 2011; Kramer and Farabaugh, 2007; Kramer et al., 2010; Zhou et al., 2015, 2009). We expect that polymorphisms that increase codon bias in domains and in 3’coding regions would be associated with faster translation rates in a population (see Table 1).

The stability of folding for mRNA secondary structures (mF) broadly influences the processing, translation, and decay of mRNA (Andrzejewska et al., 2020; Bevilacqua et al., 2016). Ribosomes transiently unwind mRNA secondary structures so codons can be read in single-stranded form (Mustoe et al., 2018; Takyar et al., 2005). Greater mF has been associated with longer ribosome pausing in vitro (Wen et al., 2008) and lower translation efficiency in bacteria (Burkhardt et al., 2017). It thus came as a surprise when it was discovered that across genes in yeast, mF is positively correlated with protein abundance (R = 0.68 from (Tuller et al., 2011; Zur and Tuller, 2012)), and appears to be selected for in highly expressed genes (Park et al., 2013).

The positive association between mF and protein abundance is not well understood but several mechanistic models have been proposed to explain how mF can both cause longer ribosome pausing and greater protein expression. Based on their simulation of yeast translation, Mao and colleagues suggest that the first few ribosomes to translate an mRNA move slowly as they unwind the secondary structures, and if those ribosomes are sufficiently slowed by the structures, then initiation rates will allow for subsequent ribosomes to pack in behind, preventing the mRNA from refolding (Mao et al., 2014). Once the mRNA is linearized and occupied by a high density of ribosomes, then relatively high quantities of protein can be produced. However, if mRNA secondary structure is weak, then elongating ribosomes proceed before subsequent ribosomes catch up, allowing the mRNA to refold between ribosomes. This results in overall slower-moving and more spaced-out ribosomes because each one must unfold the mRNA as it goes, lowering translation rates. Additionally, Zur and Tuller propose that high mF mRNAs are less prone to homodimerize and/or aggregate (Zur and Tuller, 2012). They suggest that in general, any negative effects associated with homodimerization and aggregation may well-outweigh those imparted by stable folding. Finally, Lai and colleagues observe that high mF maintains a short distance between 5’ and 3’ mRNA termini, thereby preserving favorable entropy for mRNA circularization (Lai et al., 2018). Such a looped arrangement is known to mediate translation initiation and ribosome recycling which can increase translation rates (Paek et al., 2015).

Based on the correlation of mF and protein abundance across genes and the proposed mechanistic models, we expect that polymorphisms that increase mF would be associated with higher translation rates (Table 2).

**Table 2.**
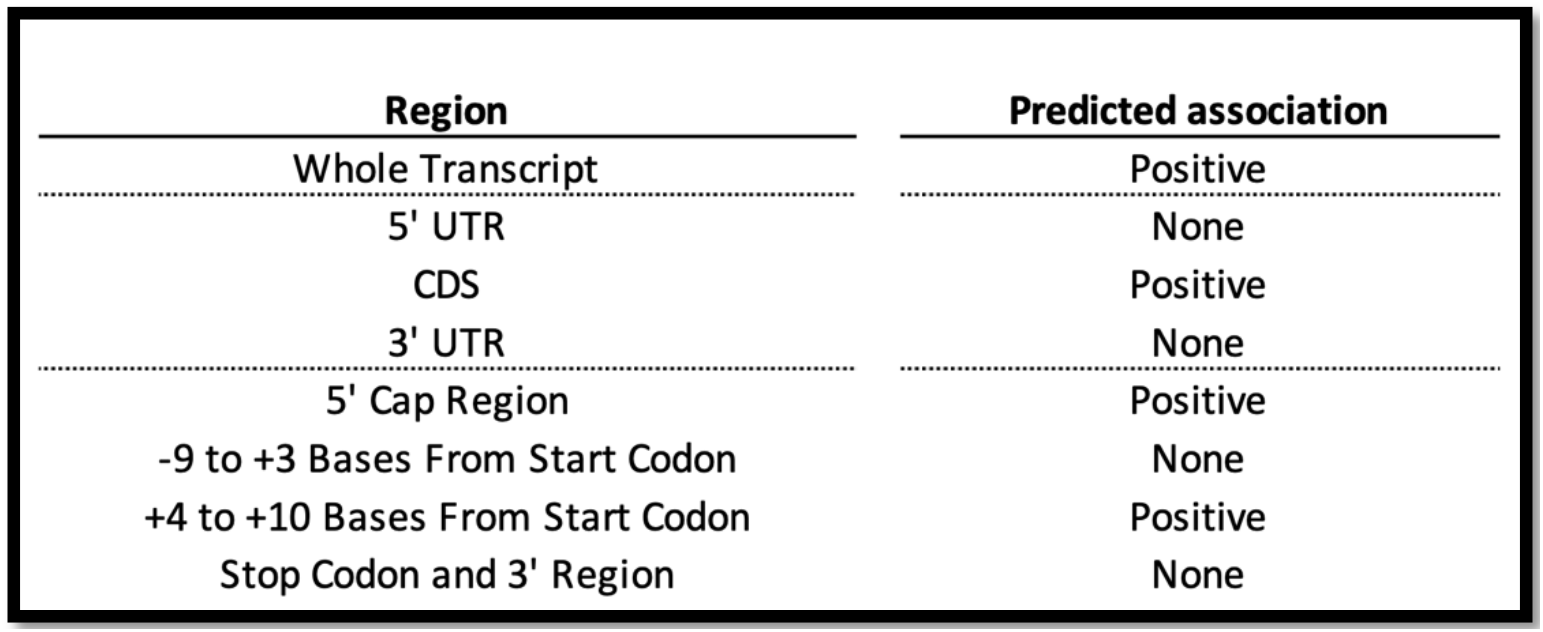
Predicted association between mF and protein expression across alleles.

It has been appreciated for several decades that stable stem-loop structures have differential impacts on protein synthesis depending on their location in an mRNA transcript (Kozak, 1986, 1989, 1990) More recent genomic approaches have revealed consensus patterns of mF across the length of mRNA transcripts and mF diversity amongst genes and taxa (Bevilacqua et al., 2016; Gebert et al., 2019).

Across all genes, the coding sequence (CDS) of yeast mRNA is more structured than either the 5’ or the 3’ untranslated region (UTR) (Kertesz et al., 2010; Wan et al., 2012). This hallmark is both selected for (Katz and Burge, 2003) and positively correlated with gene expression (Zur and Tuller, 2012). The high mF in coding sequences may boost protein expression by facilitating co-translational protein folding (Faure et al., 2016) or inhibiting unproductive translation initiation within the CDS. (Kertesz et al., 2010) We hypothesize that polymorphisms that increase mF in the CDS would therefore be associated with higher translation rates. Further, because they tend to be less structured, we hypothesize that both UTRs would show weak associations between mF and translation rates (Table 2).

In addition to the CDS and UTRs, more fine-scale regions in transcripts show mF signatures across genes and have impacts on protein synthesis. In yeast genes, high mF is associated with increased protein yield when located *+1 to +10* bases from the 5’cap (Cuperus et al., 2017; Kertesz et al., 2010). The mechanism for this association is not known and the association is in contrast with observations from mammalian mRNAs (Babendure et al., 2006; Kozak, 1989). Similarly, high mF is typically seen within the region *+4 to +10* bases from the start codon (Kertesz et al., 2010; Shabalina et al., 2006) and is hypothesized to act as a ‘speed bump’ to improve the efficiency of start codon recognition, especially in genes with suboptimal start codon contexts (Kozak, 1990). Therefore, for both the 5’ cap region and *+4* to *+10* bases from the start codon, we hypothesize that polymorphisms that increase mF would be associated with faster translation rates (Table 2).

In contrast, mF tends to be quite low within *-9* to *+3* bases from the start codon and the region from the stop codon into the 3’ UTR (Kertesz et al., 2010; Shabalina et al., 2006; Wan et al., 2012). Further, stable stem-loop structures located in these regions can inhibit translation (Kozak, 1986; Lamping et al., 2013; Niepel et al., 1999; Sherman and Baim, 1988; Vega Laso et al., 1993). We hypothesize that keeping the region *-9* to *+3* bases from the start codon and the region slightly downstream and including the stop codon free from strong mF would constrain selection for high mF across the transcript, resulting in a weak association between mF and protein synthesis rates in these regions (Table 2).

If and how codon bias and mF interact with each other to influence protein translation rates is not well understood. A simulation study (Mao et al., 2014) concluded that codon bias has the biggest impact on translation rate when mF is high because that is the scenario where ribosomes are so densely packed that the mRNA molecule becomes linearized, leaving codon bias as the rate limiting factor. Based on their results, we hypothesize that polymorphic codon bias will be most strongly associated with translation rate when mF is high.

We tested our above hypotheses by examining how allelic variation in codon bias and predicted mF each affect protein expression for 1620 genes across 22 genetically diverse *Saccharomyces cerevisiae* isolates (Skelly et al, 2013). *S. cerevisiae* is known to have strong translational selection, making this a particularly good species in which to study these factors (LaBella et al., 2019). Our findings confirm the association between codon bias and protein expression, and the association between mF and protein expression, and we extend this significance to natural variation in a single species. Most strikingly, we find that the effects of codon bias and mF are largely the consequence of their interaction, and that this interaction is more pronounced in specific regions of transcripts.

## Results

### Association of Codon Bias and Protein Expression Across 22 Yeast Isolates for 1620 Genes

To evaluate the association of codon bias and protein expression, we acquired the genome sequences, transcriptomes, and proteomes of 22 genetically diverse *S. cerevisiae* isolates sampled from six continents and 12 types of microenvironments (e.g. bee hairs, throat sputum, fermenting palm sap, leavening bread, and forest soil) (Skelly et al., 2013). Transcriptome and proteome data were measured during vegetative growth for each haploid isolate. We analyzed the 1620 genes (26.22% of 6179 total genes in *S. cerevisiae*) for which proteomic data was available in all 22 isolates. Not surprisingly, these genes are mildly enriched for housekeeping biological functions (See Methods). For each isolate’s allele of each gene, we estimated protein accumulation, independent of RNA abundance, as the natural log of the ratio of protein molecules per RNA molecule and refer to it as ‘logPPR’ (see Methods). Protein expression normalized by RNA expression is often referred to as translational efficiency and most mechanistic models connect codon bias and mF with protein synthesis rates, however, logPPR captures their effects on both protein synthesis and protein stability. We therefore refrain from using the term translational efficiency and instead use protein expression or accumulation to refer to logPPR.

Using the original measure of codon bias, the codon adaptation index (CAI) (see Methods), and logPPR, we generated a linear mixed effects regression model with logPPR as the response variable, CAI as a fixed effect explanatory variable, and gene as a random effect. By treating gene as a random effect in the mixed model, we can evaluate how allelic variation in CAI relates to logPPR for a typical gene. Over our dataset for 1620 genes, we found allelic variation in CAI to have a highly significant and positive association with logPPR (log-likelihood ratio test: G = 72.977, df = 1, *p* = 1.31e-17) (Figure 1A). Our model shows that alleles with higher codon bias tend to have higher logPPR.

**Figure 1.**
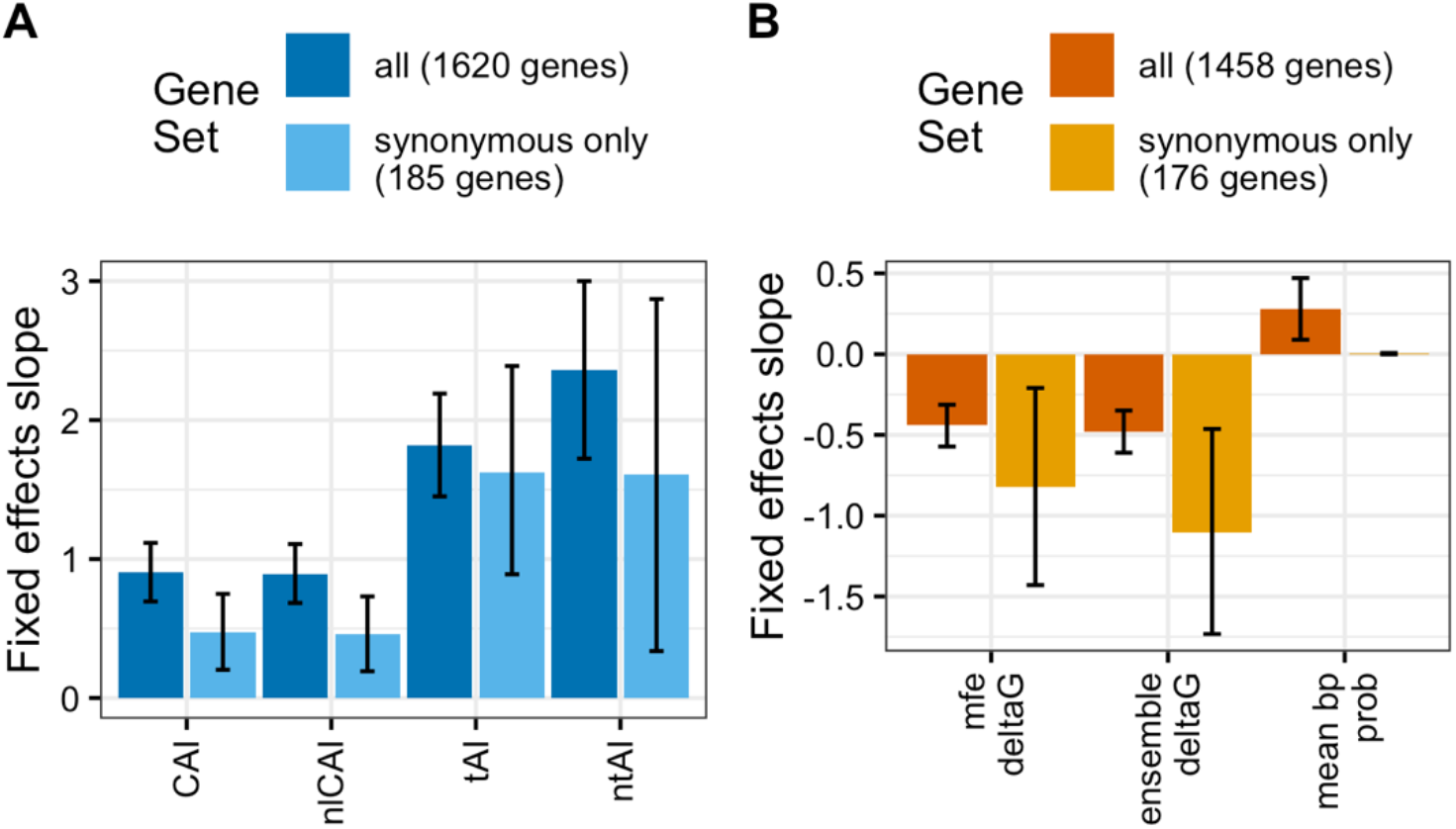
Polymorphic codon bias and mRNA secondary structure stability (mF) are each associated with protein synthesis rates. **A**, Linear mixed effects regression was used to evaluate the typical association between measures of codon bias and log protein per RNA (logPPR). Fixed effects slope of each codon bias measure (codon adaption index (CAI), length normalized codon adaptation index (nlCAI), tRNA adaptation index (tAI), normalized tRNA adaptation index (ntAI)) is shown as the predictor of logPPR. Models were computed using the full set of 1620 genes and for the 185 genes with synonymous and no non-synonymous polymorphisms. **B**, Fixed effects slope of each mF measure (minimum free energy ΔG (mfe ΔG), ensemble ΔG, and mean base-pair probability) as the predictor of logPPR in a linear mixed effects regression model. Models were computed using the full set of 1458 genes and for the 176 genes with synonymous and no non-synonymous polymorphisms. Error bars represent 95% confidence intervals.

We next examined the robustness of this result. The residuals from our model showed some heteroskedasticity (dependence on the independent variable – logPPR in this case) so we repeated our analysis using the square-root of protein molecules per RNA (sqrtPPR) as our estimate of protein accumulation. Taking the square root of a ratio is considerably less conventional than taking the log and results in a relatively compressed left tail of the distribution. This transformation eliminated the heteroskedasticity and the association between CAI and sqrtPPR was significant (log-likelihood ratio test: G = 44.135, df = 1, *p* = 3.06e-11) (Figure S1A). We note that we observed the same pattern of heteroskedasticity for logPPR and homoskedasticity for sqrtPPR for all models used throughout this study and will present logPPR results while noting differences and reporting sqrtPPR results in the supplemental figures.

Most of the genes in our study have both synonymous and non-synonymous polymorphisms. Because non-synonymous polymorphisms are known to influence logPPR through mechanisms besides codon bias, we repeated our analysis on the 185 genes that lack non-synonymous polymorphisms. Again, we found a significant and positive association between CAI and logPPR (log-likelihood ratio test: G = 11.324, df = 1, *p* = 7.65e-04) (Figures 1A & S1A).

The lengths of the 61 genes used in our CAI training set vary, such that some genes contribute more to the estimation of codon bias than others. To give each gene equal weight, we normalized codon frequencies across training set genes to calculate a normalized length CAI (nlCAI). This association between nlCAI and logPPR (log-likelihood ratio test: G = 70.982, df = 1, *p* = 3.60e-17) is negligibly different from the association between CAI and logPPR (Figures 1A & S1A).

To further evaluate if the association between codon bias and logPPR is robust to the method used to calculate codon bias, we examined two additional measures of codon bias. The tRNA Adaptation Index (tAI) measures codon bias based on tRNA gene copy number as an estimate of tRNA supply (dos Reis et al., 2003) (see Methods). The normalized tRNA Adaptation Index (ntAI) modifies tAI to also account for the demand on tRNAs by the cognate codons in the pool of mRNA (Pechmann and Frydman, 2013) (see Methods). For both our full set of 1620 genes and the synonymous-only set of 185 genes, the associations between tAI and logPPR and ntAI and logPPR are significant and positive (log-likelihood ratio tests: 1620 genes tAI G = 95.587, df = 1, *p* = 1.42e-22; 185 genes tAI G = 18.607, df = 1, *p* = 1.61e-05; 1620 genes ntAI G = 52.268, df = 1, *p* = 4.84e-13; 185 genes ntAI G = 6.1489, df = 1, *p* = 1.32e-02) (Figures 1A & S1A).

Thus, the relationship between codon bias and protein expression is robust to the method used to measure codon bias as well as to the presence or absence of non-synonymous polymorphisms. The association between tAI and logPPR using the full set of 1620 genes was the most significant of those evaluated, suggesting tRNA gene copy number is capturing the most information about the effects of codon bias on protein expression.

### Association of Polymorphic mRNA Folding Strength and Protein Expression

With the relationship between codon bias and logPPR established, we next investigated the association between mRNA folding strength (mF) and protein expression across the same 22 isolates of *S. cerevisiae*. A growing body of evidence has shown the counter-intuitive pattern that genes with more structured mRNAs produce more protein (see Introduction). For each isolate’s allele of each gene in our dataset, we calculated three measures of mF (see Methods): minimum free energy (mfe) ΔG is an estimate of the change in Gibbs free energy an mRNA experiences after folding into its most energetically stable configuration, ensemble ΔG is a Boltzmann-weighted sum of estimated ΔG values, and mean base-pair probability is the mean chance that a nucleotide is base-paired, given the weighted set of ensemble configurations. We found 1458 genes have allelic variation for these mF measures and we used these 1458 genes to evaluate the association between mF and logPPR. All three measures of mF are significantly and positively associated withlogPPR (Figures 1B & S1B).We note that because more negative ΔG represents more stable structures, we will describe a negative slope for ΔG vs logPPR as a positive association between mF and logPPR. Ensemble ΔG shows the most significant association with logPPR (log-likelihood ratio test: G = 51.861, df = 1, *p* = 5.96e-13); mfe ΔG shows nearly as significant an association (log-likelihood ratio test: G = 44.507, df = 1, *p* = 2.53e-11); mean base-pair probability shows a less significant association (log-likelihood ratio test: G = 8.2598, df = 1, *p* = 4.05e-03). To control for potential impacts of non-synonymous polymorphisms, we repeated this analysis on the 176 genes that have variation in mF and lack non-synonymous polymorphisms. We found ensemble ΔG and mfe ΔG are significantly associated with logPPR while mean base-pair probability is not (log-likelihood ratio tests: ensemble ΔG G = 11.204, df = 1, *p* = 8.16e-04; mfe ΔG G = 6.8558, df = 1, *p* = 8.84e-03; mean base-pair probability G = 0.0141, df = 1, *p* = 0.9056) (Figures 1B & S1B). Thus, we conclude that the pattern of positive association between mF and protein abundance across genes is also true for allelic variation within genes.

### Protein Expression is Predicted by an Interaction Between Polymorphic Codon Bias and mRNA Folding Strength

We next examined the interaction of polymorphic codon bias and mF. To test Mao and colleagues’ prediction that for more stable mRNA structures, codon bias plays a larger role in determining final translation elongation rates (Mao et al., 2014), we analyzed the 1447 genes polymorphic for both codon bias and mF. We used tAI to quantify codon bias and ensemble ΔG for mF because they were found to be the most significant predictors of logPPR. We computed the overall mF of a single gene as the median ensemble ΔG across alleles of the gene. We found that indeed, the top half of genes, ranked from most stable overall mF to least stable, show a much stronger relationship between polymorphic tAI and logPPR (Figures 2A & S2A). Although not a stated prediction of Mao and colleagues, for completeness we examined if the reciprocal interaction was occurring. Specifically, we wanted to determine whether highly biased genes showed a stronger relationship between mF and logPPR. We measured the overall codon bias of each gene as the median tAI across its alleles. Interestingly, we found that the top half of genes, ranked from highest overall codon bias to lowest, show a much stronger relationship between polymorphic ensemble ΔG and logPPR (Figures 2B & S2B). This pair of results suggests that codon bias and mF interact synergistically.

**Figure 2.**
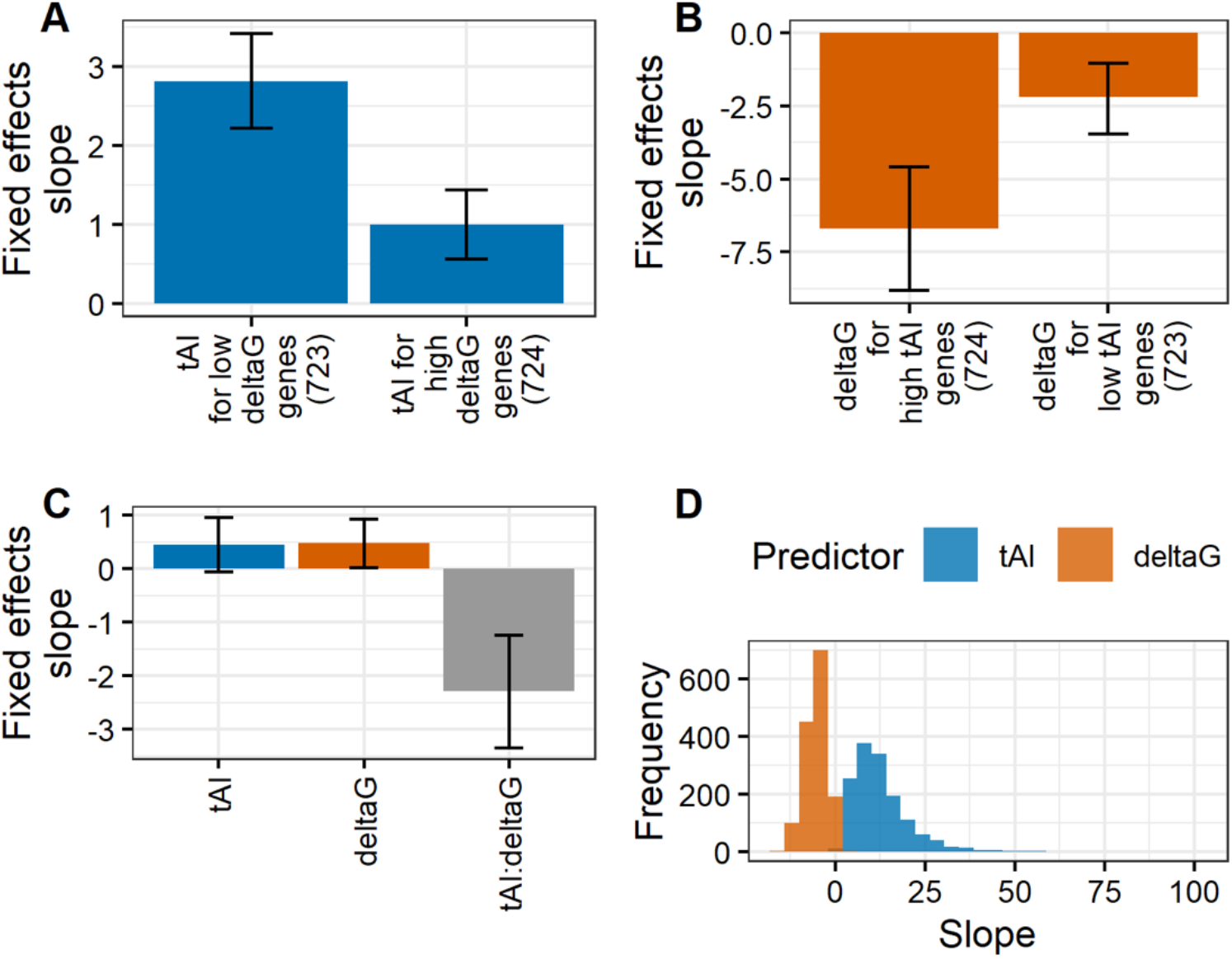
The interaction between polymorphiccodon bias and mRNA secondary structure stability (mF) is associated with protein synthesis rates. **A**, Fixed effects slope of codon bias tRNA adaptation index (tAI) as the predictor of log protein per RNA (logPPR) in a linear mixed effects regression model for the bottom and top half of genes split by median (across alleles) mF ensemble ΔG. **B**, Fixed effects slope of ensemble ΔG as the predictor of logPPR in a linear mixed effects regression model for the bottom and top half of genes split by median (across alleles) tAI. **C**, Fixed effects slope of tAI, ensemble ΔG, and tAI:ensemble ΔG interaction as the predictors of logPPR in a linear mixed effects regression model. **D**, Distribution across genes of the partial derivative of logPPR with respect to tAI and ensemble ΔG from the model with tAI, ensemble ΔG, and tAI:ensemble ΔG interaction as the predictors of logPPR. Error bars represent 95% confidence intervals.

To evaluate the interplay of individual effects and synergistic effects, we ran a linear mixed effects model with independent terms for tAI and ensemble ΔG and an interaction term between tAI and ensemble ΔG. Consistent with codon bias and mF working synergistically, the interaction term has a significant negative slope and including the interaction term significantly improves the fit of the model (log-likelihood test: G = 27.273, df = 1, *p* = 1.77e-07) (Figures 2C & S2C). Thus, stable mF and high codon bias together associate with high logPPR.

Although the term for ensemble ΔG has a weakly significant positive slope with logPPR as the independent variable, it is not significant in the model with sqrtPPR as the response variable. If increased mF inhibits protein expression, that effect is quite small compared to its effects promoting protein expression in interaction with codon bias. Indeed, the partial derivative of logPPR with respect to ensemble ΔG is negative for most genes (Figure 2D), consistent with the synergistic interaction dominating the effects.

### Role of Region-Specific Codon Bias and mRNA Folding Strength

Comparisons across genes have revealed that codon bias is strongest in domain encoding regions and in the 3’ coding regions and weakest in 5’ coding regions and inter-domain linker regions (see Introduction). As such, for the alleles of each gene, we separated those codons that fell into domain encoding and 3’ coding sequences (“domain + 3’ coding”) from those that fell into the 5’ coding and linker sequences (“5’ + linker coding”). We hypothesized that the synergistic interaction between codon bias and mF in their association with logPPR may differ between these groups. Of the 1620 genes in our dataset, 1458 have polymorphisms that alter mF. Of those, 983 have codon bias-altering polymorphisms in both domain + 3’ coding and 5’ + linker coding sequences. For these 983 genes, we ran a linear mixed effects regression model on logPPR vs. domain + 3’ coding tAI, whole transcript ΔG, and the interaction of those terms. This model confirmed that indeed, protein expression is associated with whole transcript mF and the codon bias in domain + 3’ coding sequences (Figures 3B & S3B), similar to how whole CDS tAI synergizes with whole transcript mF (Figures 2C & S3C). In contrast, the regression model of logPPR vs. 5’ + linker tAI, whole transcript ΔG, and the interaction of those terms shows no associations (Figure 3A & S3A). Thus, the interaction between codon bias and mF affects protein expression, and this is heavily driven by polymorphisms that alter codon bias in the protein domain and 3’ coding sequences.

**Figure 3.**
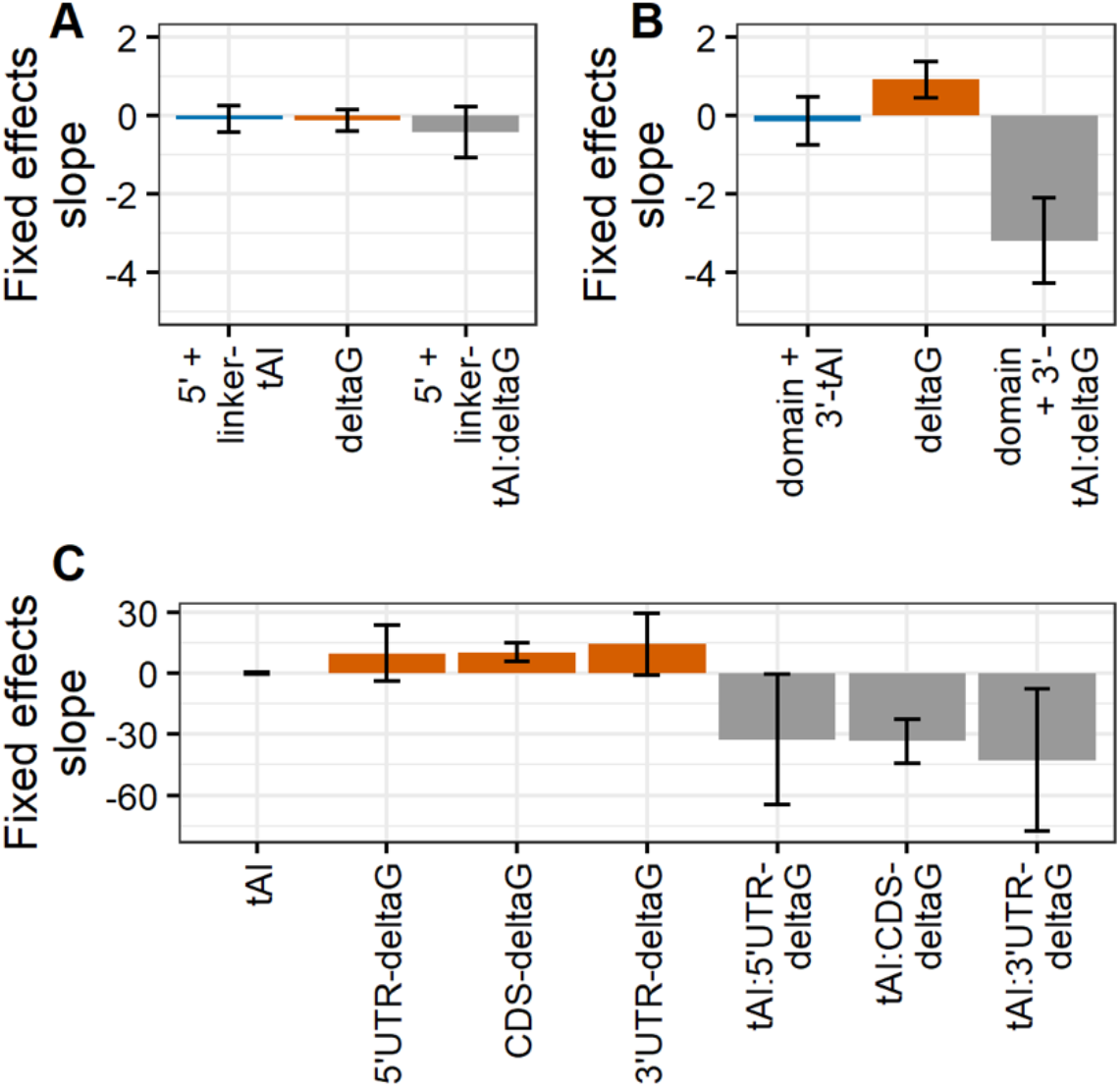
The effects of codon bias are largely due to polymorphisms localized to domain encoding and 3’ coding regions while the effects of mRNA folding stability (mF) are strongest in the CDS. **A**, To determine the localized effects of codon bias, we split coding sequences up into two regions: 5’ coding (AUG up to first domain) plus linker (sequences between domains) and domain plus 3’ coding (past last domain to stop codon). Fixed effects slope of 5’ coding plus linker region codon bias tAI, whole transcript mF ensemble ΔG, and their interaction as predictors of logPPR in a linear mixed effects regression model. **B**, Fixed effects slope of domain plus 3’ coding region tAI, whole transcript ensemble ΔG, and domain plus 3’ coding region tAI:whole transcript ensemble ΔG interaction as predictors of logPPR in a linear mixed effects regression model. **C**, To determine the localized effects of mF, we calculated proportional sum of minimum free energy (psmfe) ΔG values for substructures spanning the 5’ UTR, CDS, and 3’ UTR. Fixed effects slope of CDS tAI, 5’ UTR, CDS, and 3’ UTR mF psmfe ΔG, and CDS tAI:5’ UTR, CDS, and 3’ UTR psmfe ΔG as predictors of logPPR in a linear mixed effects regression model. Error bars represent 95% confidence intervals.

Similar to codon bias, mF varies across transcript regions (see Introduction). This led us to hypothesize that allelic differences in mF may have different effects on protein expression depending on which region’s mF they affect. We first examined the fine-scale differences in mF effects between the regions at the 5’ cap (+1 to +10 bases of 5’ cap), upstream and including the start codon (−9 to +3 bases of translation start), downstream of the start codon (+4 to +10 bases of translation start), and downstream of the stop codon (+1 to +18 bases of translation stop). In contrast with how polymorphisms act on codon bias, polymorphisms can act across a transcript to influence the mF of a distant region. Therefore, instead of categorizing polymorphisms based on their location, we looked to see how mF in each region changes across alleles. To do this we used a proportional sum of the minimum free energy (psmfe) ΔG values for individual substructures spanning a region to estimate the local mF (see Methods). For all four regions, we uncovered no significant associations between logPPR and the interaction of codon bias and mF (Figure S4A). These four regions are all quite small (<18 bp) so we looked to see if any small (40 bp) regions have significant associations and found none (Figures S4B-D), suggesting a lack of power at this scale. Next, we looked at the course-scale differences in mF effects between CDS, 5’ UTR, and 3’ UTR. Using the 1312 genes with polymorphic CDS tAI and polymorphic CDS, 5’ UtR, and 3’ U1R psmfe ΔG, we ran a linear mixed-effects model with logPPR as a function of CDS tAI, psmfe ΔG for CDS, 5’ UTR, and 3’ UTR, and the interactions between tAI and each ΔG term. This revealed that the interaction between codon bias and mF as well as the independent effects of mF on protein expression are strongest in the CDS and are weaker in the UTRs (Figures 3C & S3C). This pattern mirrors previous observations that mF tends to be higher in CDS regions relative to UTRs (Kertesz et al., 2010; Wan et al., 2012).

## Discussion

In this study, we investigated the association of allelic variation in codon bias and mRNA folding strength (mF) with allelic variation in protein expression in *S. cerevisiae*. We leveraged a published dataset of genome sequences, transcriptome abundances, and proteome abundances for 22 yeast isolates (Skelly et al., 2013), calculated codon bias and mF from genome sequence data and measured protein expression as the log of the ratio of protein levels to mRNA levels (logPPR). We removed the potential allelic effects of codon bias and mF on RNA levels and stability by focusing on the amount of protein per RNA molecule.

By using linear mixed effects models, we estimated the expected slope of the response of logPPR as a function of allelic variation in codon bias and/or mF while controlling for gene-to-gene differences in levels and effects. Although linear mixed effects models are generally robust to the assumption of homoscedasticity (model fit is consistent across values of the independent variable), logPPR did show some heteroscedasticity (model fit was better for larger values of logPPR). Reanalysis using square-root of protein per RNA (sqrtPPR) demonstrated that our results are nearly all robust to the assumption of homoscedasticity (Figures S1–3).

Previous work on codon bias and mF showed that they are each correlated with protein levels, selected for across species, and capable of altering protein levels when manipulated (Babendure et al., 2006; Dana and Tuller, 2014; Gooch et al., 2008; Hanson and Coller, 2018; LaBella et al., 2019; Mao et al., 2014; Park et al., 2013; Tuller et al., 2011; Zur and Tuller, 2012). Our study shows the most comprehensive evidence to date that allelic variation in codon bias and mF in a population are both significantly associated with the amount of protein per RNA produced (Figures 1 & S1). These associations in the context of previous work motivate a deeper investigation of codon bias and mF as important *cis*-acting mechanisms of protein expression variation.

Our findings on codon bias agree with previous studies for how codon bias alone acts on protein expression. We found that tAI, which is solely based on tRNA supply estimated from tRNA gene copy numbers, had the most significant association with logPPR (Figures 1A & S1A). Other measures of codon bias (CAI, nlCAI, and ntAI) which incorporate genomic usage of codons, were also significantly associated with logPPR, though to a lesser extent than tAI. This implies that tRNA supply is the most important aspect of codon bias in *S. cerevisiae*.

Our refined understanding of the mechanisms by which codon bias acts alone on protein expression is in sharp contrast with our speculative understanding of how mF has the counterintuitive relationship of more stable structures associating with higher protein production (Zur and Tuller, 2012). We are aware of three possible mechanistic models to explain this counterintuitive association: RNA homodimerization/aggregation avoidance, ribosome recycling via RNA circularization, and RNA structure refolding avoidance (see Introduction). Evidence exists that each could play a role, however systematic evidence is lacking.

Our result that polymorphic mF is indeed positively associated with logPPR (Figure 1B) was an important confirmation of the relationship of mF alone with protein levels. However, our examination of the interaction between codon bias and mF reframes the question about the mechanism of mF. We found that codon bias and mF act synergistically in their positive association with logPPR, that codon bias has no significant independent effects, and the independent effects of mF are negative (positive slope for ensemble ΔG vs logPPR) but only weakly significant and inconsistently so between logPPR and sqrtPPR models (Figures 2, 3, S2, & S3). Thus, the question of the mechanism of mF is more specifically a question about the mechanism by which codon bias and mF synergistically act to promote protein expression. This question remains unresolved. We have tRNA supply as an explanation for codon bias alone being positively associated with protein production and we have several possible models for mF being positively associated with protein production. However, we lack mechanistic models that explain strong synergy between codon bias and mF – strong enough that codon bias and mF have little to no independent effects.

It is noteworthy that the RNA structure refolding avoidance model described by Mao and colleagues (Mao et al., 2014) is the only model we are aware of that explicitly predicts an interaction between codon bias and mF. Their simulations concluded that codon bias is expected to have a larger effect on protein synthesis rates when mF is high but do not predict that mF has larger effects when codon bias is high. Specifically, they predict that codon bias becomes the most important factor when mF is large enough to result in a high density of ribosomes that prevents RNA secondary structure from reforming between adjacent ribosomes. Furthermore, they predict that mF at the 3’ end of transcripts would result in the biggest interaction between codon bias and mF. Although we did observe that codon bias has a larger effect when mF is high (Figures 2B & S2B), our results differed from Mao and colleagues’ in that we found mF has a larger effect when codon bias is high (Figures 2B & S2B), that the interaction between codon bias and mF is bidirectional (Figures 2C & S2C), and the regional effect of mF is highest in the CDS, not the 3’ end of the transcript (Figures 3C & S3C). Our findings suggest that either codon bias or mF could play the role of the rate limiting factor on protein expression. They also imply additional complexity in the role mF plays across the transcript than what was assumed in Mao and colleagues’ simulations. Our study will hopefully motivate future work in this area.

## Methods

### Data Collection and Processing

From the supplemental files associated with Skelly and colleagues’ manuscript (Skelly et al., 2013), we downloaded genome sequence, mRNA abundance, and peptide abundance data for the following 22 yeast isolates: 273614N, 378604X, BC187, DBVPG1106, DBVPG1373, DBVPG6765, L_1374, NCYC361, SK1, UWOPS05_217_3, UWOPS05_227_2, UWOPS83_787_3, UWOPS87_2421, Y12, Y55, YJM975, YJM978, YJM981, YPS128, YPS406, YS2, and YS9. These abundance data span the set of 1636 genes across isolates.

For each gene in each strain, we expressed protein abundance as a sum of peptide levels (Michael J. MacCoss, personal communication, July 2018); we defined the coding sequence (CDS) based on coordinates supplied by Skelly and colleagues’ supplemental general feature format (.gff) file (Skelly et al., 2013); and we defined 5’UTR and 3’UTR sequences based on UTR length specifications from Tuller and colleagues’ supplemental file (Tuller et al., 2009). The whole mRNA sequence was then the concatenation of 5’UTR, CDS, and 3’UTR sequences.

### Measuring Protein Expression with logPPR and sqrtPPR

Gene-by-gene in every strain, we measured protein expression as the steady-state ratio of protein abundance to mRNA abundance (protein per mRNA, or PPR). Before we calculated this ratio, for each isolate, we normalized mRNA abundance and protein abundance measurements by estimates of actual cell-wide mRNA and protein molecule counts (von der Haar, 2008; Miura et al., 2008). After this normalization step, rather than PPR being in arbitrary units, it is approximately in units of protein molecules per mRNA molecule. After computing PPR, we log transformed it or square root transformed it.

### Approximating Global Codon Bias with CAI, nlCAI, tAI, and ntAI

Three classic methods of estimating codon bias are the Codon Adaptation Index (CAI), the tRNA Adaptation Index (tAI), and the normalized tRNA Adaptation Index (ntAI). Each relies on its own respective codon table, where every codon maps to one value in the range (0, 1]. A gene’s CAI, tAI, or ntAI equals the geometric mean of values assigned to its comprising codons by the requisite table.

CAI quantifies a gene’s tendency to use the synonymous codons most favored by a pre-defined training set of genes (Sharp and Li, 1987). A CAI value of 1 indicates total usage of these codons, while a CAI value approaching 0 indicates complete avoidance. One approach to selecting a training set of genes is to select an arbitrary number of highly expressed genes that are presumed to reflect the strongest codon bias in the genome (Sharp et al., 1988). Ranking all genes with mRNA abundance data by their median transcript abundance (across isolates) we systematically investigated how codon usage changes as a function of selecting the 2^i^ highest expressed genes (where I ∈ [1, 12]) (Figure S5). The three sets with the largest number (1024-4096) of genes showed high frequencies of A/T rich codons, consistent with the two-fold mutational bias for A/T nucleotides over G/C nucleotides in *S. cerevisiae* (Lynch et al., 2008). The nine sets with the smallest number (2-512) of genes showed usage of codons consistent with tRNA supplies for all amino acids except cysteine and glycine. A systematic approach to choosing a training set involves algorithmically identifying the dominant codon usage bias in the genome, independent of any expression information (Carbone et al., 2003; Sharp et al., 1988). The training set of 61 genes identified by the Carbone et al. algorithm for *S. cerevisiae* has codon usage similar to the most highly expressed genes and is consistent with tRNA supplies (Figure S2). We used this training set to calculate one CAI codon table per isolate. We then computed a single median CAI codon table across isolates. This is the table we use to measure the CAI of the coding sequence (CDS) of each gene.

Normalized-by-length CAI (nlCAI) is our slightly modified version of CAI. Longer training set genes contribute more to the CAI codon table, and because all genes have their own intrinsic biases (Quax et al., 2015), these large contributions may misrepresent the dominant genomic level codon bias. Instead of computing the CAI codon table based on each gene’s synonymous codon counts, we compute it based on each gene’s synonymous codon percent abundances. Specifically, we calculate the fraction of codons that are codon *i* in each gene, and add up all such fractions across genes. This gives a 61-element array, where each value matches to a sense codon. For each group of synonymous codons, we divide their corresponding array values by the maximum array value within that group. In this way, we compute a single nlCAI codon table for each isolate, and then take their median table for nlCAI calculations.

tAI estimates how often a gene uses synonymous codons with high supplies of cognate/near-cognate tRNAs (dos Reis et al., 2003). A gene always using such codons has a tAI near 1, and a gene never using such codons has a tAI near 0. This measure accounts for cases in which one tRNA recognizes more than one codon (wobble) (Crick, 1966), and it approximates tRNA supply by tRNA gene copy number in the genome (dos Reis et al., 2004). The high positive correlation (r = 0.76) between tRNA gene copy number and tRNA abundance (in yeast) suggests that this is a reasonable approximation for our study (Tuller et al., 2010). Based on the approach by dos Reis colleagues, we compute a single tAI codon table and use it for tAI calculations in all strains (dos Reis et al., 2003).

ntAI considers both the abundance of tRNAs (as measured by tRNA gene copy number) and the abundance of codons competing for them (as measured by the sum of codon translation frequencies across all mRNAs) (Pechmann and Frydman, 2013). From this view, a codon optimal for fast translation is one whose tRNA species are high in abundance and low in demand. A gene always using such synonymous codons has a ntAI value near 1, while a gene never using such values has a ntAI value near 0. We use the Pechmann & Frydman approach (Pechmann and Frydman, 2013) to calculate an individual ntAI codon table per isolate. For each isolate, we compute ntAI with the isolate’s corresponding table.

Each measure was computed with Python (version 3.7.1).

### Approximating Local Codon Bias with tAI

We downloaded domain coordinates, as predicted by Pfam, from the *Saccharomyces* Genome Database (SGD) (date of access: February, 2019). For each gene in each isolate, we concatenated the sequences encoding Pfam-defined protein domains with the 3’ coding region (i.e. the region downstream of the 3’-most domain-encoding sequence and upstream of the translation stop codon). This is the “domain+3’ coding” mRNA region. For the “linker+5’ coding” mRNA region, we concatenated the sequences encoding any inter-domain linkers with the 5’ coding sequence (i.e. the region downstream of the start codon and upstream of the 5’most domain-encoding sequence). Using Python (version 3.7.1), we then computed tAI, our chosen measure of codon bias, for domain + 3’coding and linker + 5’coding regions.

### Approximating Global mRNA Folding with Mean Base-Pair Probability, mfe ΔG, and Ensemble ΔG

Three gauges of mRNA folding are mean base-pair probability, minimum free energy (mfe) ΔG, and thermodynamic ensemble ΔG. All are predicted for entire mRNA transcripts (at 30°C) with the RNAfold algorithm (version 2.4.14) from the ViennaRNA Package (Lorenz et al., 2011).

Mean base-pair probability is the arithmetic mean of nucleotide pairing probabilities. One such pairing probability represents the chance that a given nucleotide is in a base-paired configuration, given the weighted set of thermodynamic ensemble configurations. It is calculated via the partition function (McCaskill, 1990). A mean base-pair probability near 1 suggests that an mRNA’s folded form is highly structured and stable.

Minimum free energy (mfe) ΔG represents the change in Gibbs free energy an mRNA experiences after folding into its most energetically stable (mfe) configuration, as predicted by RNAfold. A negative ΔG value of large magnitude indicates spontaneous formation of a highly stable structure.

Ensemble ΔG is a Boltzmann-weighted sum of ΔG values; one ΔG value per mRNA structure in the mRNA’s thermodynamic ensemble. Because mfe structure is only a best-guess prediction and because mRNA folding is far from static (Crothers et al., 1974), ensemble ΔG is expected to be a more accurate measure of overall mRNA folding strength.

### Proportional Sum Mean Free-Energy ΔG

To calculate mF for regions within a transcript, we used the RNAeval tool from the ViennaRNA Package (version 2.4.14) (Lorenz et al., 2011) we obtained a detailed thermodynamic description of each gene’s mfe structure at 30°C. Specifically, the algorithm reports a ΔG approximation for all substructures that fully describe an mRNA’s overall folding shape: multi loops, external loops, interior loops, and hairpin loops. To compute the ΔG of an mRNA region (e.g. the CDS), we first summed the ΔGs of all substructures completely enclosed within it. Then, for any partially enclosed substructure, we 1) calculated what fraction of the substructure is built by nucleotides from our region, 2) multiplied this value by the substructure’s ΔG, and 3) added the result to our existing sum. We called this value the proportional sum mean free-energy (psmfe) ΔG.

### Gene Criteria and GO Term Enrichment Analyses

Limitations in the availability of data and which genes contained variation across isolates for the explanatory variables in our models required us to compute our models with different sets of genes. Here, we explain how these gene sets were selected and we summarize the results of their Gene Ontology (GO) term enrichment analyses.

i. Of the 1636 genes with mRNA and protein abundance data across isolates, 1620 show one or more SNPs across isolates. Of these, 185 show only synonymous SNPs. To obtain the latter information, we translated the 1636 coding sequences from each isolate via the translate tool from the SeqIO Biopython package (Cock et al., 2009). For each gene, we then aligned the corresponding set of amino acid sequences (one sequence from each isolate) via the MUltiple Sequence Comparison by Log-Expectation (MUSCLE) algorithm (Edgar, 2004). Those genes with 100% amino acid identity scores and SNP(s) across isolates were used in our 185 gene analyses. In considering model results based on this smaller set of genes, we were able to discount any effect amino acid substitutions may have on translation rates.
ii. We used 1458 of 1620 genes in our models of global mF. These genes have available length data for the 5’UTR and the 3’UTR, and they have one or more SNPs in their concatenated 5’UTR, CDS, and 3’UTR sequences. Of these 1458, 176 have 100% amino acid identity for our synonymous gene set.
iii. The intersection of the 1620-gene set and the 1458-gene set defines the set of 1447 genes we used in our analyses of the synchronous actions of codon bias and mF. We ranked these 1447 genes by their median tAI across isolates, chose the bottom 723 genes as our ‘low tAI’ group and the top 724 genes as our ‘high tAI’ group. This process is repeated for ensemble ΔG in place of tAI.
iv. In the models pertaining to regional codon bias, we considered a 983 gene subset of the 1447 genes defined above. Each gene belonging to this subset is characterized by an absence of premature stop codons, available protein domain region prediction data from Pfam, and SNP(s) in both the domain + 3’ coding sequence and the linker + 5’coding sequence.
v. To arrive at a subset of genes suitable for regional structure models, we filtered the 1447-gene set defined above by the following criteria to generate an 779-gene set: genes must have variation (across isolates) in local mfe ΔG within the 5’UTR, the CDS, the 3’UTR, *+1 to* + *10* from the 5’cap, *-9 to +3* from translation start, *+4 to +10* of translation start, and *+ 1 to +18* from translation stop. Additional criteria were 5’UTRs at least 19 nucleotides in length and 3’UTRs at least one nucleotide in length.

With few exceptions, our GO-term enrichment analyses show that genes in every set are most enriched for GO-terms related to 1) general metabolism, 2) nucleotide synthesis and metabolism (purine’s especially), 3) peptide biosynthesis and metabolism, 4) amino acid synthesis and metabolism, 5) ATP metabolism, and 6) translation. This result was not unexpected as all isolates were grown at a steady-state temperature of 30°C in nutrient rich broth, they were all sampled at log-phase growth, and mass spectrometry most reliably detects highly expressed proteins. GO-term enrichment results were generated by the PANTHER overrepresentation test (released April, 2020) via the GO biological process complete annotation for *S. cerevisiae* (version 2020-03-23).

### The Linear Mixed Effects Regression Model

We computed all linear mixed effects regression models and log-likelihood ratio tests with the lme4 package (version 1.1.21; Bates et al, 2015) from R (version 3.6.0). Each computed model has one explanatory variable with ‘gene’ as the random effect (both slope and intercept).

### Data Availability

Data files and analysis scripts are available at https://github.com/anastacia9/bias_mF.

## Acknowledgements

We thank Mike MacCoss for guidance on approximating protein abundances and Aidan Corbin, Olivia Dong, Benjamin Haagen, Suzanne Lee, Dietmar Schwarz, and Tara Wirsching for helpful discussions and comments on the manuscript. This work was supported by National Science Foundation Award MCB-1518314 (D.A.P. 2015) and Western Washington University.

## Author contributions

A.W. and D.A.P. conceptualized and designed the study. A.W. analyzed the data and M.B. performed additional validation. A.W., M.B., and D.A.P. interpreted the data, generated figures, and wrote the manuscript.

## Supplemental Files

**Figure S1.**
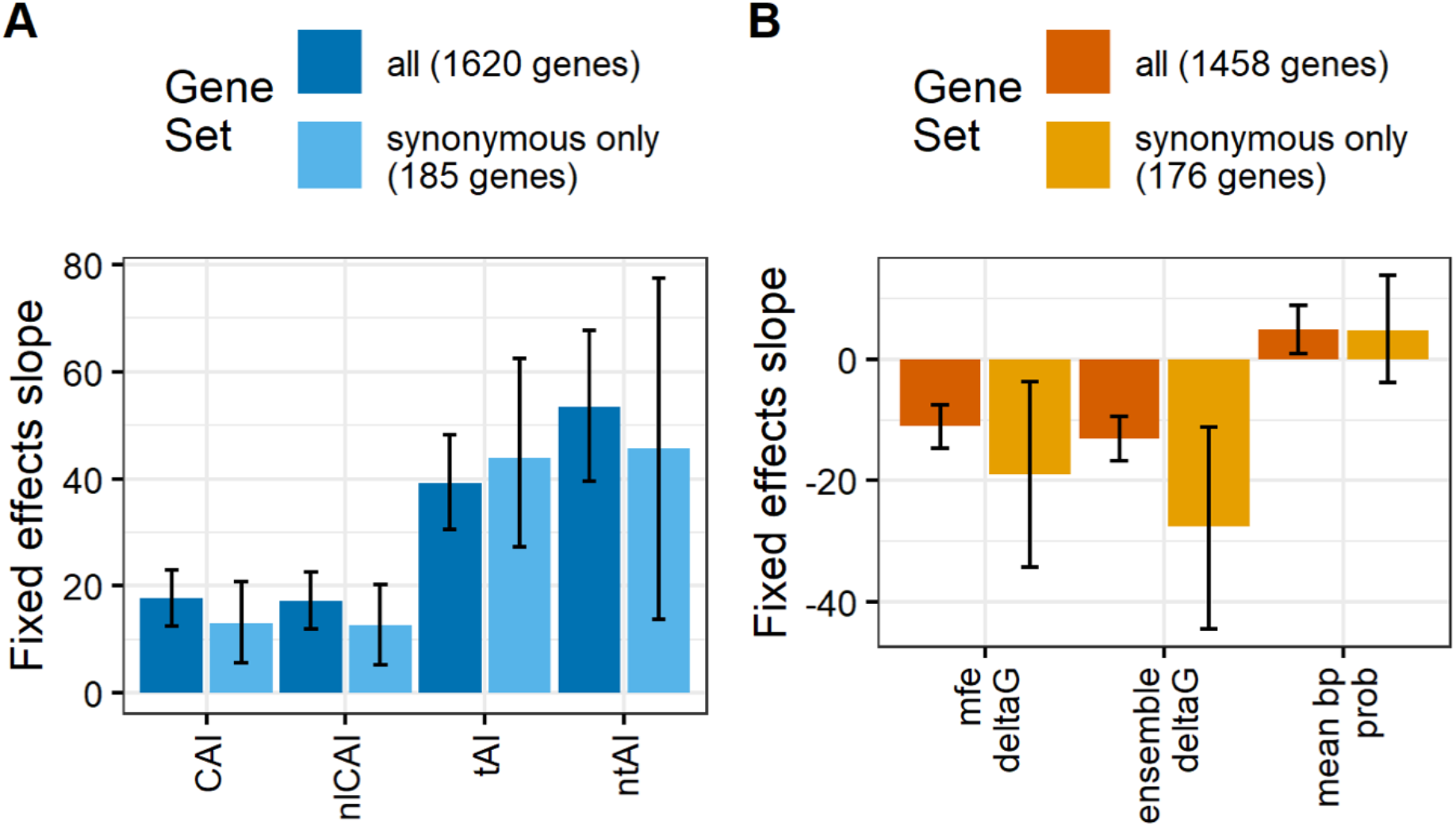
Polymorphic codon bias and mRNA secondary structure stability (mF) are each associated with protein expression as measured by the square-root of protein molecules per RNA molecule. **A**, Fixed effects slope of each codon bias measure (codon adaption index (CAI), length normalized codon adaptation index (nlCAI), tRNA adaptation index (tAI), normalized tRNA adaptation index (ntAI)) as the predictor of square root protein per RNA (sqrtPPR) in a linear mixed effects regression model. Models were computed using the full set of 1620 genes and for the 185 genes with synonymous and no non-synonymous polymorphisms. **B**, Fixed effects slope of each mF measure (minimum free energy ΔG (mfe ΔG), ensemble ΔG, and mean base-pair probability) as the predictor of logPPR in a linear mixed effects regression model. Models were computed using the full set of 1458 genes and for the 176 genes with synonymous and no non-synonymous polymorphisms. Error bars represent 95% confidence intervals.

**Figure S2.**
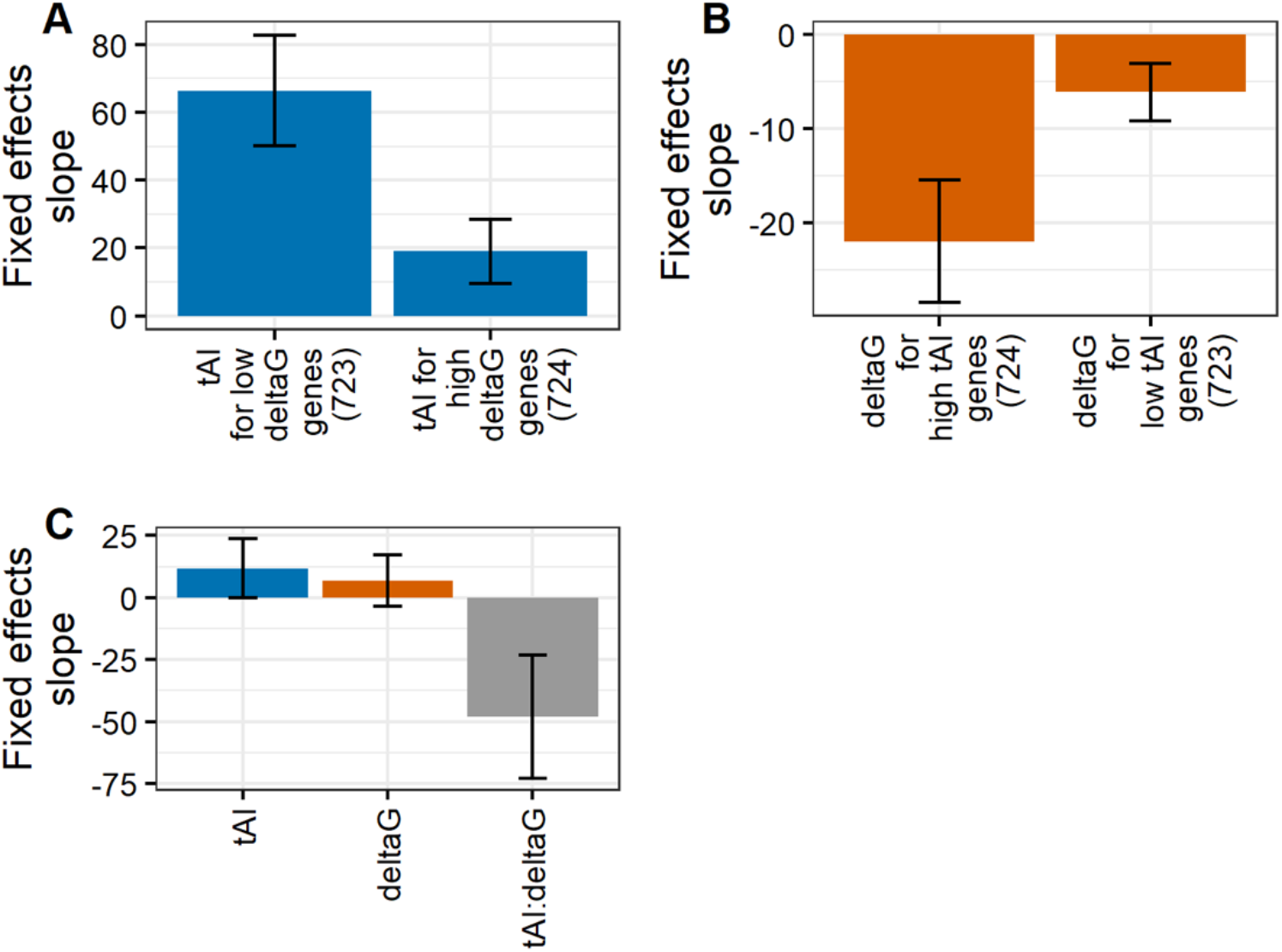
Polymorphic codon bias and mRNA secondary structure stability (mF) interact in association with protein expression as measured by the square-root of protein molecules per RNA molecule. **A**, Fixed effects slope of codon bias tRNA adaptation index (tAI) as the predictor of square root protein per RNA (sqrtPPR) in a linear mixed effects regression model for the bottom and top half of genes split by median (across alleles) mF ensemble ΔG. **B**, Fixed effects slope of ensemble ΔG as the predictor of sqrtPPR in a linear mixed effects regression model for the bottom and top half of genes split by median (across alleles) tAI. **C**, Fixed effects slope of tAI, ensemble ΔG, and tAI:ensemble ΔG interaction as the predictors of sqrtPPR in a linear mixed effects regression model. Error bars represent 95% confidence intervals.

**Figure S3.**
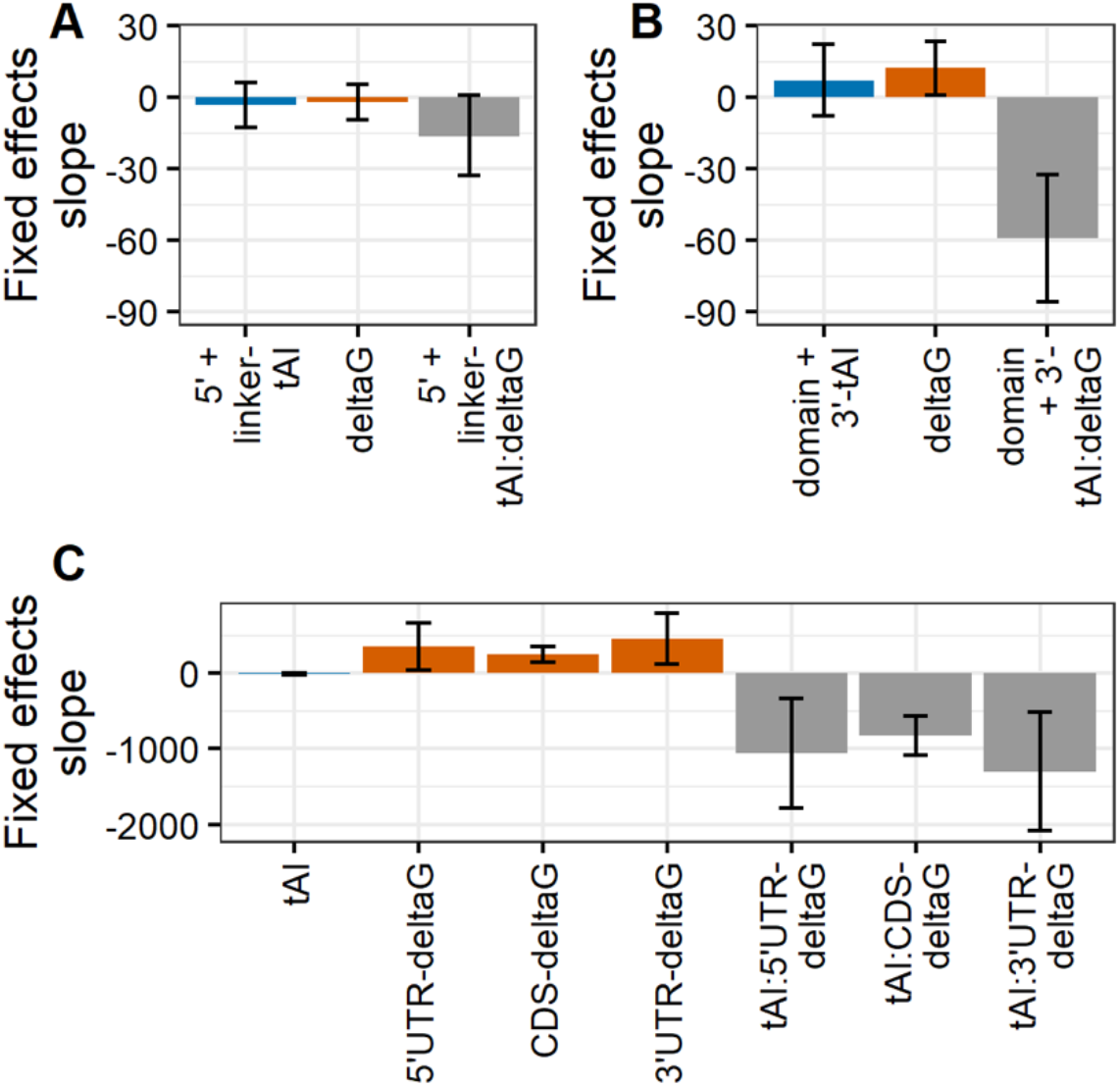
The effects of codon bias on square root protein molecules per RNA molecule (sqrtPPR) are largely due to polymorphisms localized to domain encoding and 3’ coding regions while the effects of mRNA folding stability (mF) on sqrtPPR are strongest in the CDS. To determine the localized effects of codon bias, we split coding sequences up into two regions: 5’ coding (AUG up to first domain) plus linker (sequences between domains) and domain plus 3’ coding (past last domain to stop codon). **A**, Fixed effects slope of 5’ coding plus linker region codon bias tAI, whole transcript mF ensemble ΔG, and 5’ coding plus linker region tAI:whole transcript ensemble ΔG interaction as predictors of square root protein molecules per RNA molecule (sqrtPPR) in a linear mixed effects regression model. **B**, Fixed effects slope of domain plus 3’ coding region tAI, whole transcript ensemble ΔG, and domain plus 3’ coding region tAI:whole transcript ensemble ΔG interaction as predictors of sqrtPPR in a linear mixed effects regression model. To determine the localized effects of mF, we calculated proportional sum of minimum free energy (psmfe) ΔG values for substructures spanning the 5’ UTR, CDS, and 3’ UTR. **C**, Fixed effects slope of CDS tAI, 5’ UTR, CDS, and 3’ UTR mF psmfe ΔG, and CDS tAI:5’ UTR, CDS, and 3’ UTR psmfe ΔG as predictors of sqrtPPR in a linear mixed effects regression model. Error bars represent 95% confidence intervals.

**Figure S4.**
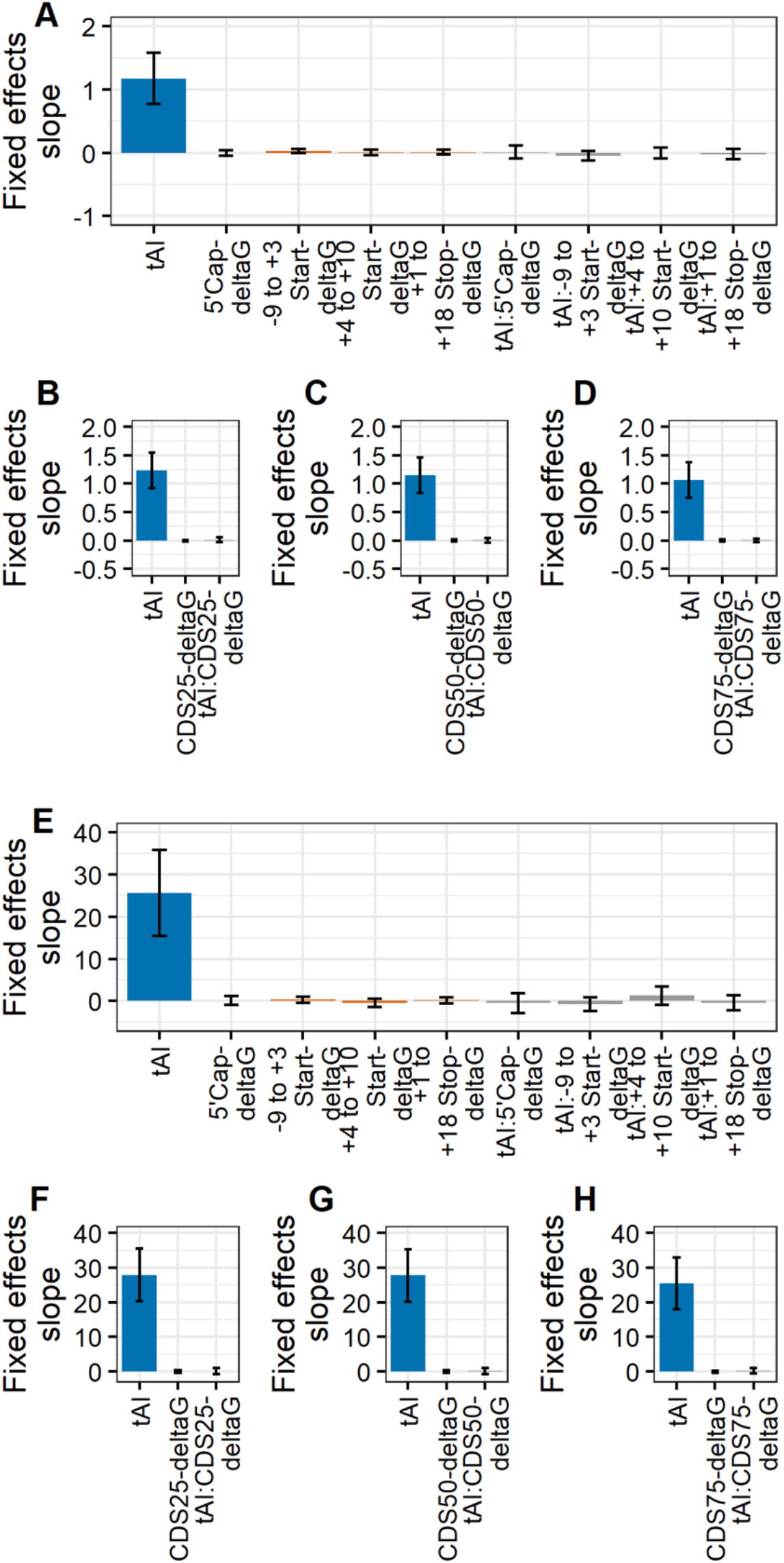
No evidence of fine-scale effects of mRNA folding stability (mF) on protein synthesis. To determine the fine-scale localized effects of mF, we calculated proportional sum of minimum free energy (psmfe) ΔG values for substructures spanning the 5’ cap (+1 to +10), just before and including the start codon (−9 to +3), just after the start codon (+4 to +10), and just after and including the stop codon (+1 to +18). **A & E**, Fixed effects slope of CDS tAI, 5’ cap, −9 to +3 start, +4 to +10 start, and +1 to +10 stop mF psmfe ΔG, and CDS tAI:5’ cap, −9 to +3 start, +4 to +10 start, and +1 to +10 stop mF psmfe ΔG as predictors of logPPR (A) or sqrtPPR (E) in a linear mixed effects regression model. To evaluate our power to detect smallscale effects we sampled 40 bp regions located at 25%, 50%, and 75% of the total CDS length and calculated psmfe ΔG values for these regions. Fixed effects slope of CDS tAI, 25% (**B & F**), 50% (**C & G**), and 75% (**D & H**) CDS mF psmfe ΔG, and CDS tAI:25%, 50%, and 75% CDS mF psmfe ΔG as predictors of logPPR (B-D) or sqrtPPR (F-H) in a linear mixed effects regression model. Error bars represent 95% confidence intervals.

**Figure S5.**
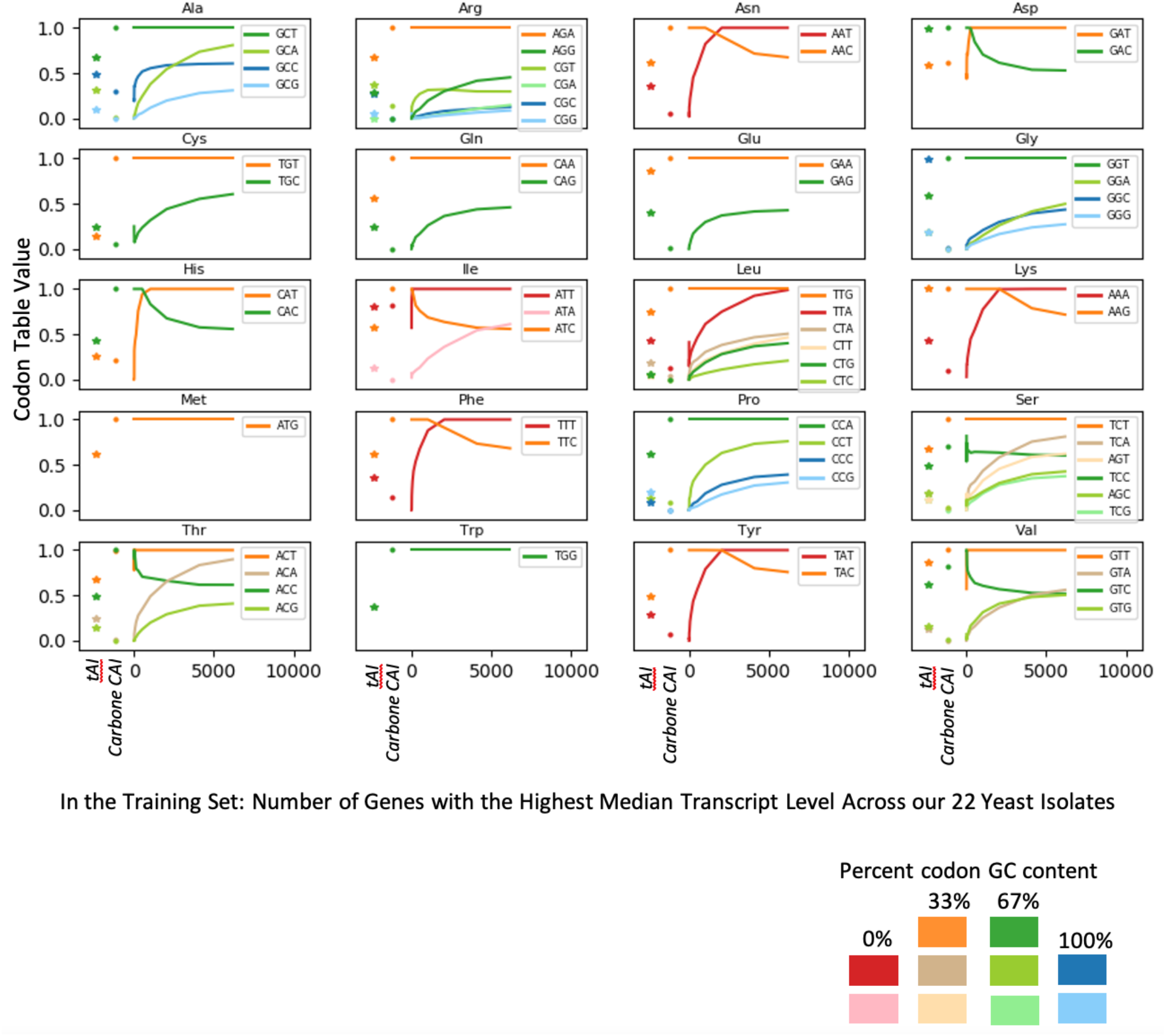
Codon Table Values. Lineplot showing how CAI codon table values change in response to the number of high expressing genes in the CAI training set. Datapoints are taken for training sets containing 6179 or 2i (where i [1,12]) of the most highly expressed genes (as ranked by median transcript abundance acros our 22 yeast isolates). For each amino acid, the most common synonymous codon among training set genes has a value of 1. A sibling synonymous codon appearing 60% as often would have a value of 0.6. Each subplot corresponds to an amino acid and its synonymous codons. Codon color is based on %GC content: red (0% GC), orange (33% GC), green (67% GC), and blue (100% GC). For reference, we also present each codon’s codon table value from Carbone and colleagues (Carbone et al., 2003) as well as each codon’s tAI codon table value.

